# Click chemistry-based quantification of extracellular matrix turnover for drug screening and regenerative medicine

**DOI:** 10.1101/2025.05.08.652928

**Authors:** Annie Porter, Songshan Fan, Ying Peng, Mengxi Lv, Yilu Zhou, Abdulaziz Alanazi, Lin Han, Liyun Wang, X. Lucas Lu

**Affiliations:** Department of Mechanical Engineering, University of Delaware, Newark, Delaware, 19716, USA; School of Biomedical Engineering, Science and Health Systems, Drexel University, Philadelphia, PA 19104, USA

**Keywords:** Click Chemistry, Metabolic labeling, Glycosaminoglycan, Collagen, Cartilage, Tissue engineering

## Abstract

This study presents a sensitive and cost-efficient method to quantify extracellular matrix (ECM) synthesis and degradation using copper-free click chemistry reactions to fluorescently label new ECM components. The approach enables spatial visualization and longitudinal measurement of specific ECM turnover *in vitro*. We validated the method across multiple platforms, including native cartilage explants and monolayer cultures of human mesenchymal stem cells and breast cancer cells. The technique also proved effective for osteoarthritis drug screening by detecting compounds that mitigate inflammation-induced ECM degradation. Compared to traditional biochemical or histological assays, this click chemistry-based technique offers higher sensitivity, lower sample requirements, and improved temporal resolution. Its versatility supports broad applications in tissue engineering, regenerative medicine, disease modeling, and high-throughput drug evaluation.

## INTRODUCTION

The extracellular matrix (ECM) of connective tissues such as cartilage, bone, and ligament is a highly organized network of proteins, proteoglycans, and other biomacromolecules. The ECM’s chemical composition, mechanical properties, and reservoir of signaling molecules are key in controlling the cell niche and cell behaviors.^1^ Healthy ECM is a dynamic structure that undergoes continuous modeling and remodeling through the metabolic activities of cells.^2^ Dysregulation of ECM turnover is associated with many diseases, such as fibrotic diseases, tumor growth, osteoporosis, and osteoarthritis.^3^ Biomedical studies of connective tissues and ECM-driven diseases often require precise measurements of ECM dynamics, such as the synthesis and degradation rates of ECM molecules like collagen and glycosaminoglycan (GAG).^3^ In such *in vitro* studies, cells are preferably cultured in their native ECM to maintain a cell phenotype close to the *in vivo* state.

Biochemical assays and antibody- and dye-based stains are commonly used to characterize the macromolecular composition of the ECM. Staining techniques visualize macromolecule spatial distributions, whereas biochemical assays quantify their content. The sensitivity of biochemical assays is a critical component of research design. In tissue engineering studies, for example, cartilage cells seeded in hydrogels often need to be cultured for weeks to generate detectable levels of new GAG or collagen measured by dimethylmethylene blue [DMMB] (for sulfated GAG) or hydroxyproline (for collagen) assays.^4^ In addition, most staining and biochemical assay techniques offer only “snapshot” measurements, representing a one-time test following the termination of cell culture. Few of these techniques can longitudinally track ECM turnover or differentiate between newly synthesized ECM and pre-existing ECM in the tissue. Isotopic radiolabeling can quantify newly synthesized ECM macromolecules, but it lacks the spatial resolution of staining-based techniques, and its limited biocompatibility for cells makes it unsuitable for long-term tracking of ECM synthesis.^5^

Click chemistry reactions have recently been explored for labeling proteins and ECM macromolecules. Click chemistry, coined by K.B. Sharpless in 2001, represents “a set of powerful and highly reliable reactions for the rapid synthesis of new compounds and combinatorial libraries through heteroatom links.” ^6^ Click chemistry reactions are modular, high-yielding, and rapid. They produce minimal byproducts and need only simple reaction conditions.^6^ The classic click reaction, copper-catalyzed azide-alkyne cycloaddition (CuAAC), has been widely used in pharmaceutical and material science research.^7,8^ However, the copper catalyst is toxic to mammalian cells at the required reaction concentration, precluding applications in living cells. To address this limitation, a new class of click reactions was developed, termed “bioorthogonal” by C.R. Bertozzi, that can occur in a living system without interfering with the native biochemical processes.^9^ A widely used reaction is strain-promoted azide-alkyne cycloaddition (SPAAC), commonly referred to as copper-free click chemistry, which eliminates the need for a copper catalyst by employing ring strain release of cyclooctyne derivative to drive the reaction.^10^ Ligation products based on bioorthogonal click chemistry reactions have been developed for proteomic and glycobiology research.^11,12^ These tools hold significant promise for cell bioengineering or regenerative medicine research. However, they have not yet been widely adopted to evaluate ECM dynamics in biomedical studies. For example, the DMMB assay remains the most common method for quantifying GAG content in cartilage and osteoarthritis studies, and the hydroxyproline assay is still the first-line option for collagen measurement. There remains a strong demand for quantitative techniques that assess ECM turnover with high sensitivity and specificity, particularly for investigating how cell metabolism responds to environmental stimuli such as mechanical loading, growth factors, inflammation, and drug treatment.

In this study, we present a copper-free click chemistry (SPAAC)-based method to quantify ECM synthesis in monolayer and native tissue culture by fluorescently labeling nascent GAG and collagen macromolecules synthesized by cells. Benefiting from the bioorthogonality of the SPACC reaction, we further demonstrate a complementary method to quantify ECM degradation by longitudinally tracking GAG and collagen loss in tissue during long-term culture. Using these new techniques, we quantified cell metabolic activities and ECM dynamics in response to mechanical stimuli, inflammatory challenge, and drug treatment. The click chemistry-based methods proved accurate, convenient, and cost-efficient for evaluating ECM turnover in cell and tissue culture, with potential applications in biology, drug screening, personalized medicine, and regenerative medicine research.

## RESULTS AND DISCUSSION

### Metabolic Labeling of Glycosaminoglycans and Collagen via Click Chemistry

We employed strain-promoted azide-alkyne cycloaddition (SPAAC) reactions to visualize and quantify the synthesis of ECM macromolecules during short- and long-term culture. In brief, molecular probes conjugated with azide groups were added into the cell/tissue culture medium for a specific time, termed the metabolic labeling period. The molecular probes enter cells, serving as building blocks for synthesizing nascent ECM molecules (Fig. 1A Step 1). Afterwards, a dibenzocyclooctyne (DBCO)-modified fluorophore was conjugated to the azide groups on the new ECM molecules through the cycloaddition click reaction (Fig. 1A Step 2,3).

**Figure 1.**
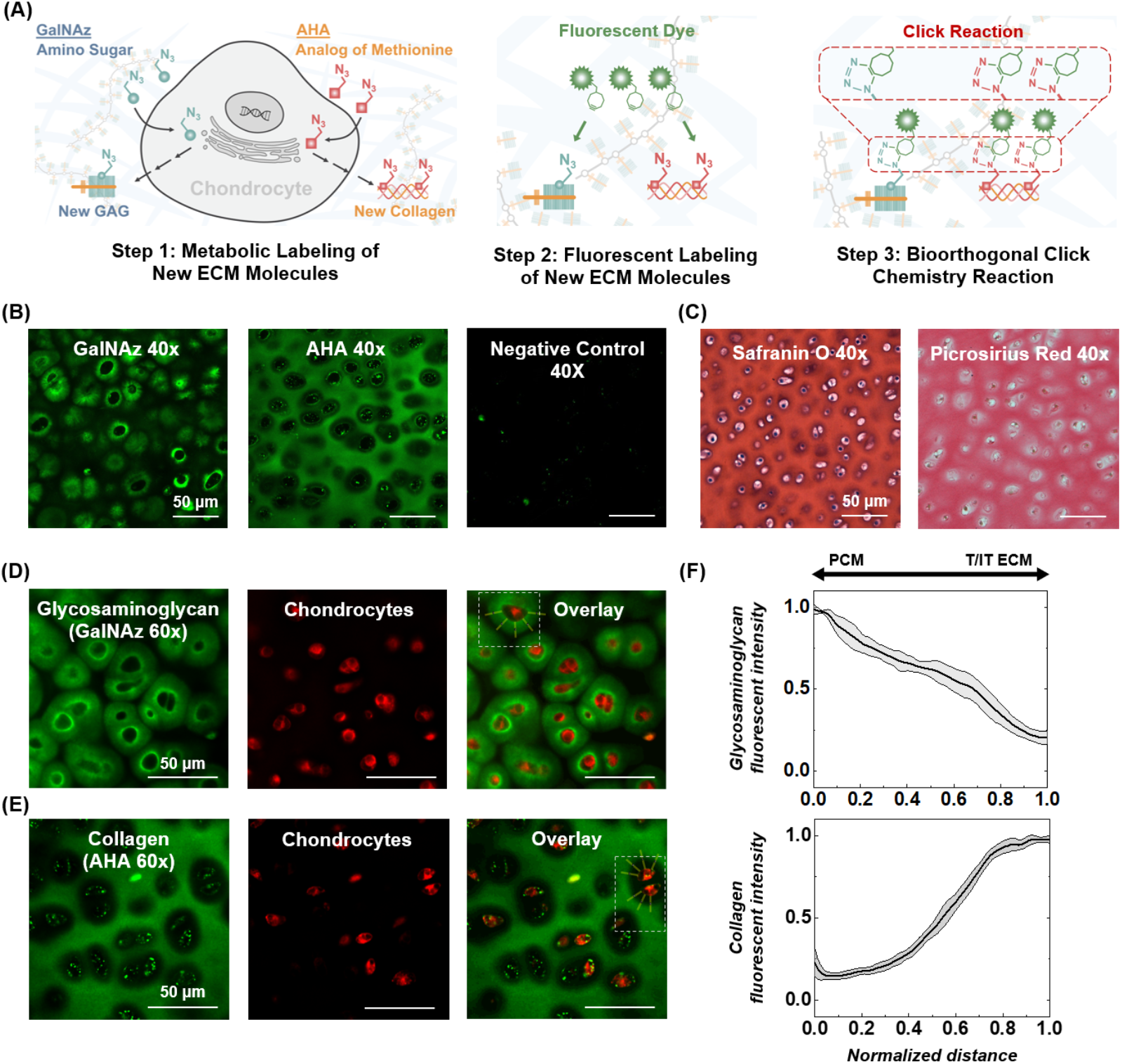
Click chemistry method to label nascent extracellular matrix (ECM) molecules in cartilage. (A) Labeling protocol. Step 1: Azide-modified molecular probes are incorporated by cells into newly synthesized ECM components. GalNAz is incorporated into glycans (left side of cell schematic), and AHA into proteins (right side of cell schematic). Step 2,3: A dibenzocyclooctyne (DBCO)-modified fluorescent dye “clicks” to the azide-labeled matrix molecules in a copper-free click chemistry reaction. The nascent ECM molecules are fluorescent for further imaging or quantification. (B) Representative confocal images (40X magnification) of calf articular cartilage labeled with GalNAz or AHA using the click chemistry method. Negative control received the same treatment without metabolic labeling. (C) Histological staining of total sulfated glycosaminoglycan content (Safranin O) and collagen content (Picrosirius Red) in bovine calf cartilage. Confocal images (60X magnification) of click chemistry labeled (D) glycosaminoglycans or (E) collagen (green) overlaid with chondrocytes (CellTracker™ Red CMPTX). (F) Representative spatial distributions of glycosaminoglycan and collagen fluorescent intensity ±1 SD were measured from the fluorescent images. Fluorescent intensity was evaluated with increasing radial distance from the cell membrane, from the pericellular matrix (PCM) of the cell into the territorial-ECM (T-ECM) and inter-territorial ECM (IT-ECM) of the cartilage.

To label newly synthesized glycans (sugar chains) with the click chemistry method, N-azidoacetylgalactosamine-tetraacylated (GalNAz),^12,13^ an azide-modified N-acetyl-galactosamine (GalNAc), was added to the cell or tissue culture medium. GalNAz can be converted by the cell GalNAc salvage pathways into the nucleotide sugar UDP-GalNAz, incorporated into O-linked glycans, and further into the disaccharide building blocks of chondroitin sulfate GAG chains.^14,15^ Similarly, L-Azidohomoalanine (AHA), an azido-analog of the amino acid methionine, was incorporated during metabolic labeling to label newly synthesized proteins, analogous to traditional radiolabeling methods in cartilage using ^35^S-labeled methionine.^11,16^ AHA is accepted as a substrate by methionyl-tRNA synthetase,^11^ and can be used to synthesize new proteins without altering cellular processes.^5,17–19^ Following metabolic labeling, the azide-labeled glycans or proteins were conjugated with the fluorescent dye AZ488 DBCO through the SPAAC click reaction.^20^ The reaction is bioorthogonal, requiring no toxic catalysts and producing no toxic byproducts. In this study, each sample was labeled using only one probe, GalNAz or AHA, but not both as they are detected with the same fluorophore AZ488 DBCO. After the click reaction, newly synthesized glycans or proteins are fluorescent, enabling visualization and quantification (Fig. 1B).

We first used the click chemistry method to visualize the spatial distribution of newly synthesized glycans and proteins in articular cartilage. Fresh cartilage harvested from bovine calf knee joints was metabolically labeled in chondrogenic medium supplemented with either GalNAz or AHA for 24 hours, followed by fluorescent labeling with AZ488 DBCO for 4 hours.^21^ After washing off excess dye, cartilage samples were imaged using confocal microscopy (Fig. 1B). The newly synthesized GalNAz labeled glycans localized around each cartilage cell (chondrocyte), forming a fluorescent halo with intensity decreasing radially from the plasma membrane (Fig. 1D). Overlap with cell-tracker labeled images suggested that newly synthesized glycans were attached on the plasma membrane. In contrast, AHA-labeled nascent proteins were dispersed throughout the cartilage’s interterritorial space of ECM (IT-ECM).^22^ Fluorescence was minimal in the pericellular ECM (PCM) and territorial ECM (T-ECM) regions, creating a distinct dark area immediately surrounding chondrocytes (Fig 1D).

Proteoglycans and collagen constitute over 95% of the ECM synthesized by chondrocytes, with collagen accounting for ∼60% and proteoglycans ∼35%.^23^ The most abundant proteoglycan in cartilage is aggrecan, which contains up to 100 attached chondroitin sulfate GAG chains.^15,24^ These chains consist of alternating amino sugar GalNAc and glucuronic acid units.^25^ Thus, the supplied azide-modified GalNAz can be incorporated into the newly synthesized chondroitin sulfate chains. Cartilage contains three distinct pools of aggrecan in the ECM: the pericellular, territorial, and interterritorial regions.^26^ Our click chemistry labeling confirms that the nascent aggrecans preferentially localize within the PCM region, consistent with the known behavior of new GAGs attaching to CD44 on the chondrocyte surface.^26–29^ They formed a bright ring ∼5 μm thick around the plasma membrane, surrounded by a halo of diminishing intensity extending an additional 5-15 μm outward (Fig. 1F). For context, mature chondrocytes in calf cartilage have a diameter of ∼10 µm, and a PCM thickness of ∼4-7 µm^30,31^.

Cartilage collagen molecules contain higher methionine than other ECM proteins, such as the aggrecan core protein (see Supplementary Table 1), and are more abundant throughout the cartilage ECM. Thus, most AHA molecules are expected to incorporate into collagen. Here, click chemistry labeling revealed minimal nascent collagen in the PCM, in contrast to nascent GAG (Fig. 1E,F). Collagen molecules rapidly migrated to the IT-ECM within 24 hours of secretion by chondrocytes, where they may be assembled into new collagen fibrils or integrated into the existing collagen network.^32^ A similar pattern was noted in 3D hydrogels seeded with mesenchymal stem cells, in which newly deposited collagen predominantly resided in the IT-ECM, leaving the PCM virtually devoid of collagen.^33^

We further compared the fluorescent images of nascent GAG and collagen in cartilage with traditional dye-based histological staining. Safranin O, a cationic dye, binds to the negatively charged sulfated GAGs, and picrosirius red, an anionic dye, has a high affinity for cationic collagen fibers.^34,35^ Unlike click chemistry, which label only newly synthesized ECM, histological dyes stain the total GAG or collagen content of the ECM. Nevertheless, Safranin O revealed higher staining intensity around the chondrocytes, whereas picrosirius red staining was more intense throughout the IT-ECM region but weaker in the PCM/T-ECM regions (Fig. 1C). Thus, histological stains revealed that areas with highly accumulated nascent GAG and collagen also exhibited higher over GAG and collagen contents, respectively.

### Quantifying ECM Synthesis Rate

In addition to visualizing the newly synthesized ECM in native tissue, click chemistry can quantify ECM synthesis rates with high sensitivity and accuracy. After labeling cartilage samples using the click chemistry method, we enzymatically digested the tissue and measured fluorescent intensity of the resulting solution using a fluorescent plate reader (Fig. 2A). To assess the feasibility and accuracy of this method, we examined the synthesis rate of *in situ* chondrocytes within cartilage under inflammatory stimulation by interleukin-1β (IL-1β), a key proinflammatory cytokine in joint inflammation that disrupts cellular metabolism and contributes to cartilage degradation.^20,36^ Cartilage samples from calf knee joints were cultured with GalNAz or AHA for 24 hours during IL-1β exposure, either without IL-1β pre-treatment (0 Pre + 1 Day) or after seven days of IL-1β pre-treatment (7 Pre +1 Day) (Fig. 2B,C). Exposure to 1 ng/ml IL-1β for 24 hours reduced the amount of nascent GAG (0.77±0.04 vs 1.00±0.02, p<0.05), but did not significantly affect collagen synthesis (0.9±0.2 vs 1.0±0.3, p>0.05) (Fig. 2B,C). A 7-day pre-treatment with IL-1β significantly reduced the 24-hour synthesis of both GAG and collagen.

**Figure 2.**
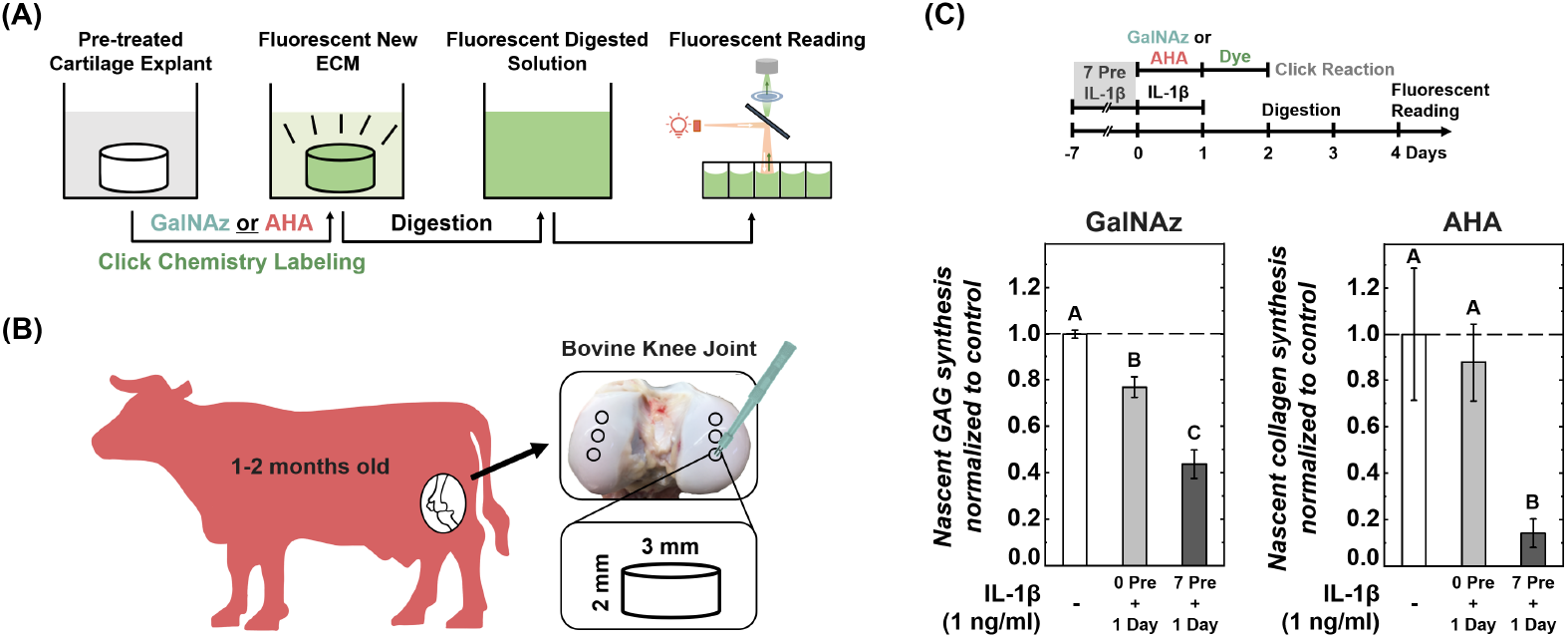
Quantifying ECM synthesis rates in articular cartilage with the click chemistry method. (A) Newly synthesized glycosaminoglycan (GAG) or collagen was fluorescently labeled with GalNAz or AHA, respectively, using the click chemistry method. The labeled samples were digested enzymatically, and the fluorescence of the digested solution was quantified using a plate reader. (B) Articular cartilage samples were harvested from bovine calf knee joints. (C) Synthesis of nascent GAG and collagen in 24 hours. Synthesis was quantified during simultaneous 24-hour exposure to pro-inflammatory cytokine IL-1β (1 ng/ml), either without (0 Pre + 1 Day) or with (7 Pre + 1 Day) a 7-day pre-treatment of IL-1β. Fluorescent readings were normalized to sample weight and then to control (n = 10, n = no. of samples per group). Different letters indicate significant differences between groups (p<0.05) tested via one-way ANOVA with Tukey post hoc analysis. Error bars represent ±1 SD.

Nascent GAG synthesis was decreased by another ∼40% compared to the 0 Pre + 1 Day group (0.44±0.06, p<0.05), and collagen synthesis reduced by ∼85% compared to the 0 Pre + 1 Day group (0.14±0.06, p<0.05). As little as 3 mg of cartilage would be sufficient to detect the synthesis differences within a 24-hour culture period under inflammatory conditions. This sensitivity contrasts with traditional methods such as the DMMB GAG assay, which typically requires longer culture durations (weeks) and larger tissue samples to detect cytokine effects on GAG synthesis.^37^ Since the click chemistry method shortens the culture period necessary to quantify nascent ECM deposition, it could optimize artificial tissue fabrication in response to parameters including cell source, cell density, scaffold selection, growth factors, and physical stimuli.^38,39^ Additionally, due to its minimal tissue requirements, the method holds potential in oncology for optimizing therapeutic strategies and dosages, making it suitable for use with tumor biopsy samples.^40,41^

### Quantifying ECM Synthesis of Cells Cultured in Monolayer

Human bone marrow-derived mesenchymal stem cells (hMSCs), widely used in regenerative medicine for treating musculoskeletal diseases,^42^ share similarities with chondrocytes in predominantly producing collagens and glycans. Here, hMSCs were seeded at ∼10,000 cells per well in a 96-well plate and cultured in chondrogenic differentiation media.^43^ After 24 hours of attachment, GalNAz or AHA was added to the culture medium for 12, 24, 36, 48, 60, or 72 hours. Azide-labeled nascent ECM molecules were then tagged with AZ488 DBCO (Fig. 3A). After 48 hours, both nascent glycans and collagens were clearly visible around the hMSCs (red) (Fig. 3B). Glycans from adjacent cells merged to form networks, unlike the new collagen molecules, which remained isolated around each cell. This pattern differed from that observed in native cartilage matrix (Fig. 1C, D). Enzymatic digestion showed no increase in fluorescent intensity during the first 24 hours, consistent with the initial ECM synthesis inactivity of seeded hMSCs as reported in the literature.^44^ Significant ECM synthesis was detected starting at 36 hours, with further increases every 12 hours thereafter (Fig. 3C) (p<0.05). Here, the click chemistry method proved sensitive enough to quantify ECM deposition by as few as 10,000 hMSCs within 12-hour increments, proving it a powerful tool for tissue engineering and regenerative medicine applications.

**Figure 3.**
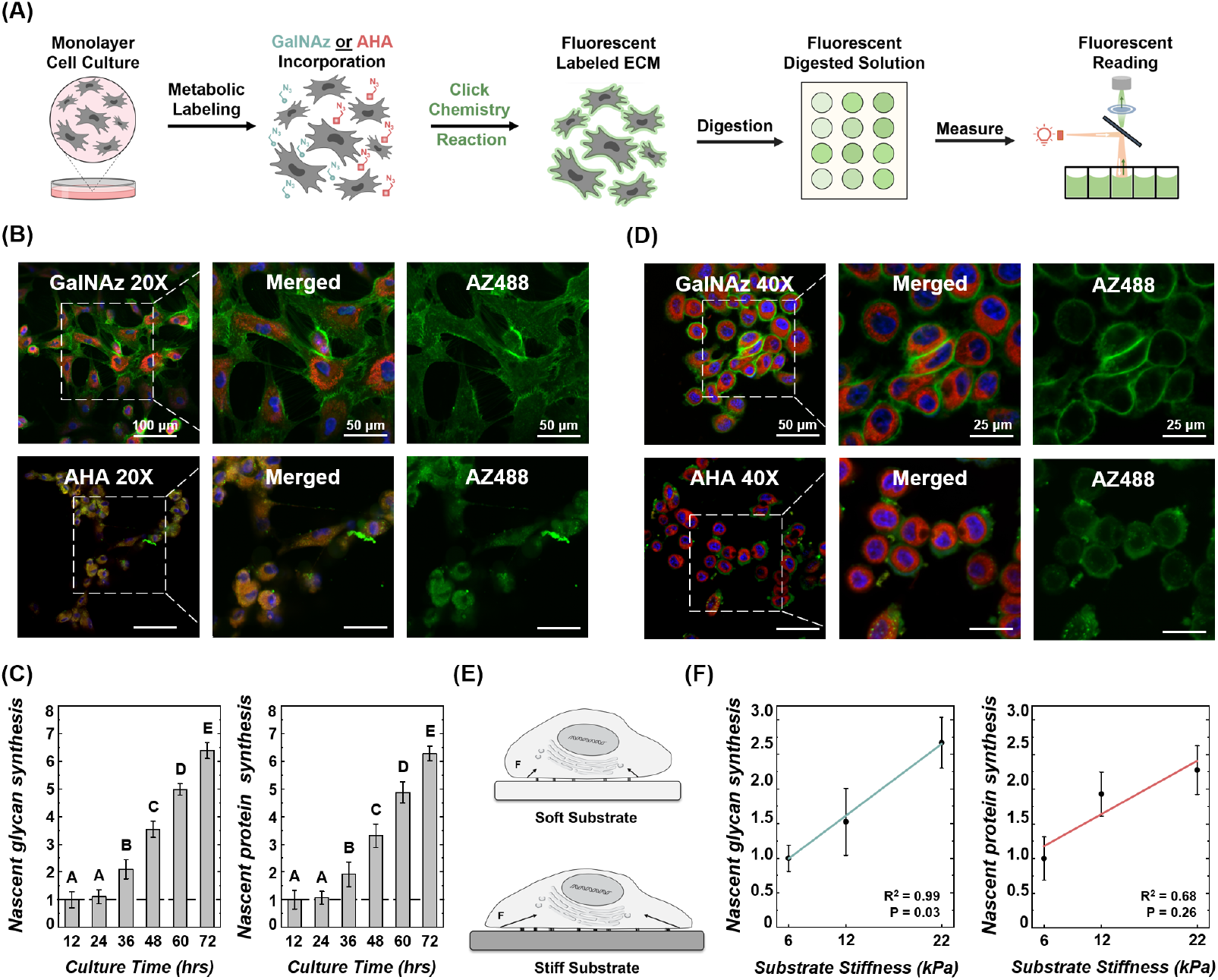
Quantifying ECM deposition of stem cells and cancer cells cultured in monolayer. (A) Nascent glycan or protein synthesis by human MSCs was metabolically labeled with GalNAz or AHA, then fluorescently labeled through the click chemistry reaction. Cells and nascent ECM were enzymatically digested, and the fluorescence of the digestion solution was quantified on a plate reader. (B) Nascent glycan (GalNAz) and protein (AHA) deposition (green), synthesized by hMSCs (cell: CellTracker™ Red CMPTX, nuclei: Hoechst 33342) in 48 hrs, imaged on a confocal microscope (40X magnification). (C) Synthesis of nascent glycans and proteins by 10,000 hMSCs in 12-72 hours (n = 10, n = no. of replicates). Synthesis rates at all time points were normalized to the initial 12-hour reading. Different letters indicate significant differences between groups (p<0.05) tested via one-way ANOVA with Tukey post hoc analysis. (D) Nascent glycan (GalNAz) and protein (AHA) deposition (green) by SKBR3 cancer cells (cell: CellTracker™ Red CMPTX, nuclei: Hoechst 33342) in 48 hrs, imaged on a confocal microscope (40X magnification). (E) Schematic of cells on substrates with varying stiffness. (F) Synthesis of nascent glycans and proteins by 10,000 SKBR3 breast cancer cells on substrates of different stiffness after 36 hours of metabolic labeling with GalNAz or AHA (n = 10). Synthesis rates were normalized to the 6 kPa value. Correlation of ECM synthesis rates with substrate stiffness assessed by simple linear regression. Error bars represent ±1 SD.

Using the click chemistry method, we investigated how substrate stiffness affected ECM deposition in breast cancer cells,^45^ given the ECM’s critical role in tumor remodeling, cancer progression, and metastasis.^46–49^ Increased ECM stiffness has been reported to promote malignant phenotypes.^46,50^ SKBR3 breast cancer cells were first cultured on a glass slide and metabolically labeled with GalNAz or AHA for 48 hours. The nascent glycans and proteins were predominantly localized around the cells, with glycans forming a thinner halo closer to the plasma membrane compared to the methionine-containing proteins (Fig. 3D). We then cultured SKBR3 cells on polyacrylamide films of various stiffness (6, 12, and 22 kPa) (Fig. 3E).^51^ Cells were metabolically labeled with GalNAz or AHA for 36 hours followed by dye labeling and enzymatic digestion. Consistent with previous studies, ECM synthesis rates increased with substrate stiffness,^52,53^ with the lowest synthesis on 6 kPa substrates and the highest on 22 kPa substrates (Fig 3F). The click chemistry method captured the characteristics of ECM deposition by stem cells and cancer cells, demonstrating its potential across multiple cell types.

### Tracking Degradation of Cartilage ECM

Copper-free click chemistry reactions are biorthogonal, allowing for longitudinal tracking of cartilage degradation without disrupting tissue culture. During long-term culture, fluorescent-labeled ECM molecules may degrade and diffuse into the culture medium. By measuring the fluorescent intensity in the medium, we can quantify the degradation of click-labeled ECM molecules from the same sample over time (Fig. 4A). Here, nascent GAG was metabolically labeled for 24 hours using click chemistry and then monitored for longitudinal loss of GAG from cartilage under inflammatory conditions. IL-1β was added to the culture medium to promote chondrocyte catabolic activities and cartilage degradation.^20^ Labeled cartilage was cultured in chondrogenic medium supplemented with 0.25, 0.5, or 1 ng/ml IL-1β for 8 days, with the medium replaced every other day. The fluorescence of the spent culture medium was measured. After 8 days, cartilage samples were enzymatically digested to assess the remaining labeled GAG in the tissue. The cumulative GAG loss was normalized to the total fluorescence of the culture medium and the tissue digest (Fig. 4A). Using a similar protocol, collagen loss from cartilage samples was tracked every three days for 18 days.

**Figure 4.**
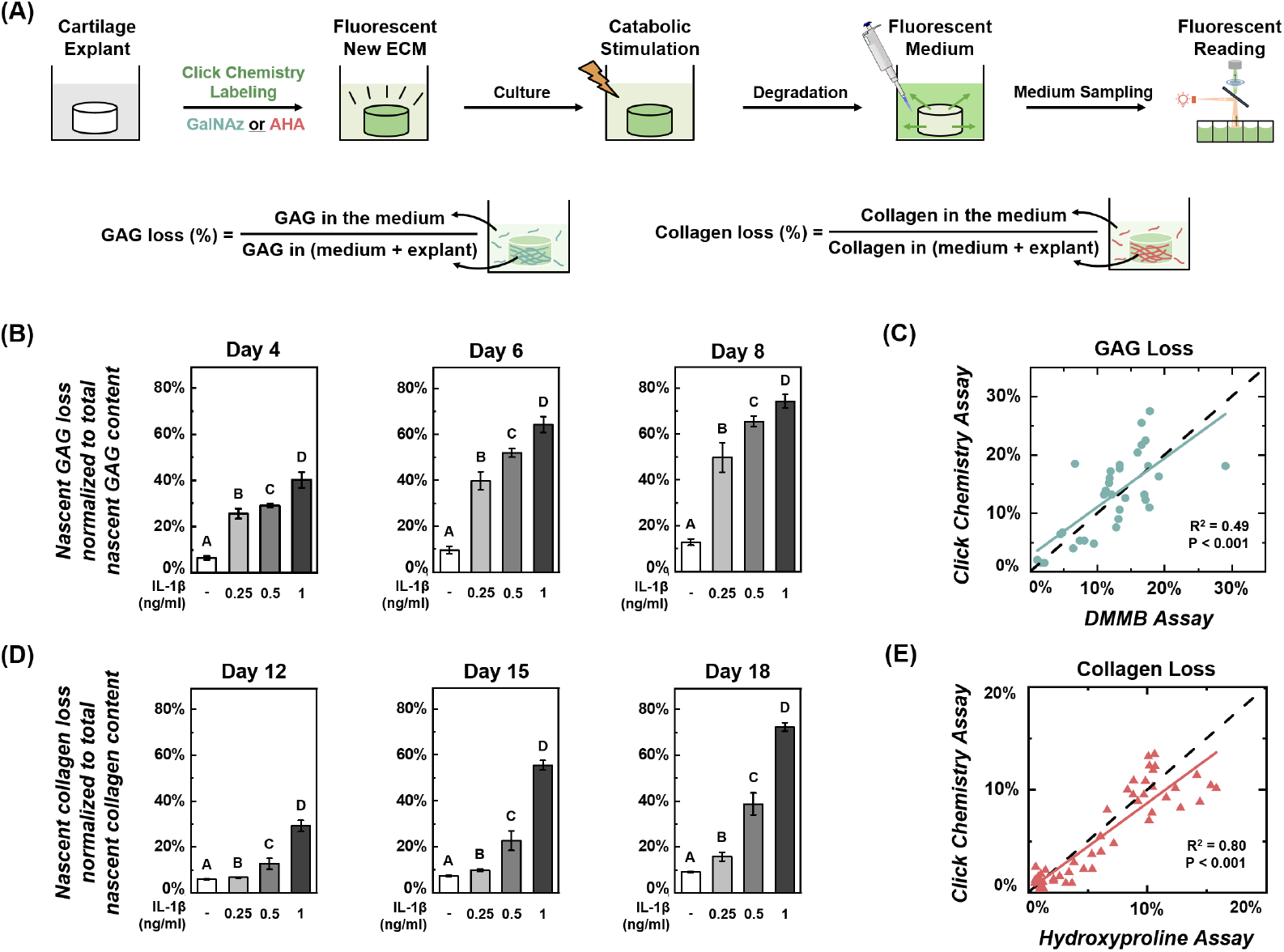
Longitudinal tracking of ECM degradation in cartilage during long-term culture. (A) Experimental protocol for tracking longitudinal ECM degradation during tissue culture. Cartilage ECM components, GAG or collagen, were first labeled using the click chemistry method. During the following culture, ECM degradation was induced by an inflammatory cytokine, causing the release of fluorescently labeled molecules, GalNAz or AHA-labeled ECM, into the culture medium. The fluorescent intensity of spent medium was read on a plate reader. The cumulative GAG or collagen loss was normalized to the total fluorescence of the culture medium and the tissue digest. (B) Cumulative GAG loss from cartilage in the presence of 0.25, 0.5, and 1 ng/ml IL-1β over an 8-day culture (n = 10, n = no. of samples per group). (C) Comparison of daily readings of GAG loss from the click chemistry and DMMB chemical assays. (D) Cumulative collagen loss from cartilage over an 18-day culture (n = 10). (E) Comparison of daily readings of collagen loss from the click chemistry and hydroxyproline assays (dashed line is y = x).

In young, healthy cartilage, the loss of labeled GAGs became evident after just 4 days of exposure to IL-1β and continued to increase during the culture period. By day 8, 0.25 ng/ml IL-1β resulted in a 50±6% total loss of nascent GAG, compared to 13±1% in control samples. GAG loss increased significantly with higher IL-1β doses, with 0.5 ng/ml causing a 65±2% total loss and 1 ng/ml inducing 74±3% loss (for all IL-1β concentrations: p<0.05 vs ctrl) (Fig. 4B). In contrast, no significant loss of the labeled collagen was detected during the first 12 days of IL-1β exposure (Fig. 4D), consistent with previous studies that showed collagen degradation begins ∼10 days later than GAG loss from *in vitro* cultured cartilage.^20,54,55^ After an 18-day IL-1β exposure, 0.25 ng/ml IL-1β induced greater collagen loss compared to control (16±2% vs 9.1±0.3%), 0.5 ng/ml IL-1β caused 39±5% total loss, and 1 ng/ml induced 72±2% total loss (for all IL-1β concentrations: p<0.05 vs ctrl). GAG and collagen loss rates were dose-dependent, indicating the influence of joint inflammation on cartilage degradation.^56^

Traditional sGAG and hydroxyproline assays were also performed using the same spent culture medium collected during the click chemistry studies. sGAG content was measured using the DMMB assay,^57^ which relies on the binding interaction between the cationic dye DMMB and the anionic sulfate groups of GAGs. A significant correlation was detected between the GAG loss rates determined by the DMMB assay and those determined using click chemistry (Fig. 4C, R^2^ = 0.49, p < 0.001). Similarly, collagen loss estimated by the hydroxyproline assay strongly correlated with the click chemistry results (Fig. 4E, R^2^ = 0.80, p<0.001). The temporal trends of GAG and collagen loss detected by the click chemistry and traditional chemical assays were highly consistent across individual samples (see Supplementary Figure S1).

The click chemistry methods offer significant advantages in terms of cost and efficiency. The chemicals required for the click chemistry method cost approximately one-fifth of those needed for traditional assays (Click: ∼$1.50/sample; DMMB assay: ∼$5/sample, and Hydroxyproline: ∼$7/sample). When tracking longitudinal ECM degradation, click chemistry becomes more cost-efficient, as losses at each time point can be quantified with simple fluorescence plate reader measurements rather than repeated biochemical assays. Thus, click chemistry represents a rapid, low-cost, and convenient tool for quantifying the ECM remodeling and kinetics during tissue culture.

### Osteoarthritis Drug Screening

To explore the potential of click chemistry for high-throughput drug screening, we evaluated the effects of three small molecule drugs (lithium chloride, valproic acid, and dexamethasone) on inflammation induced degradation of cartilage ECM, a hallmark of osteoarthritis.^58,59^ Lithium Chloride, a Wnt signaling agonist, inhibits Hedgehog signaling in chondrocytes,^60,61^ which is essential for cell metabolic behaviors. Valproic acid inhibits IL-1 induced mPGES-1 expression in chondrocytes,^62^ reducing the release of matrix metalloproteases (MMPs), key proteases mediating cartilage ECM degradation. Dexamethasone is a corticosteroid that can ease joint inflammation.^63^ We investigated the protective effects of these drugs on cartilage under inflammatory conditions. Cartilage samples were first labeled with GalNAz or AHA, and then cultured in a medium supplemented with (1) IL-1β (1ng/ml), (2) IL-1β + Lithium Chloride (LiCl, 5 mM), (3) IL-1β + Valproic acid (VPA, 5 mM), and (4) IL-1β + Dexamethasone (Dex, 100 µM). Samples were cultured for 12 days to monitor GAG loss or 28 days for collagen loss. Dexamethasone significantly reduced GAG loss compared to the IL-1β alone group, 65±6% vs 83±2% (p<0.05) (Fig. 5A). LiCl and VPA had no significant impact on GAG loss. After 28 days, IL-1β caused a substantial loss of nascent collagen (86±3%). All three drugs alleviated collagen loss significantly (p<0.05 vs IL-1β). Dexamethasone almost eliminated inflammation-induced collagen loss, bringing it to the control group levels (13±1% vs 12±2%). VPA reduced collagen loss to 25±2%, and LiCl reduced it to 73±4% (Fig. 5B). This study demonstrated that click chemistry is an effective and convenient approach for evaluating drug efficacy in preventing ECM degradation under inflammatory conditions.

**Figure 5.**
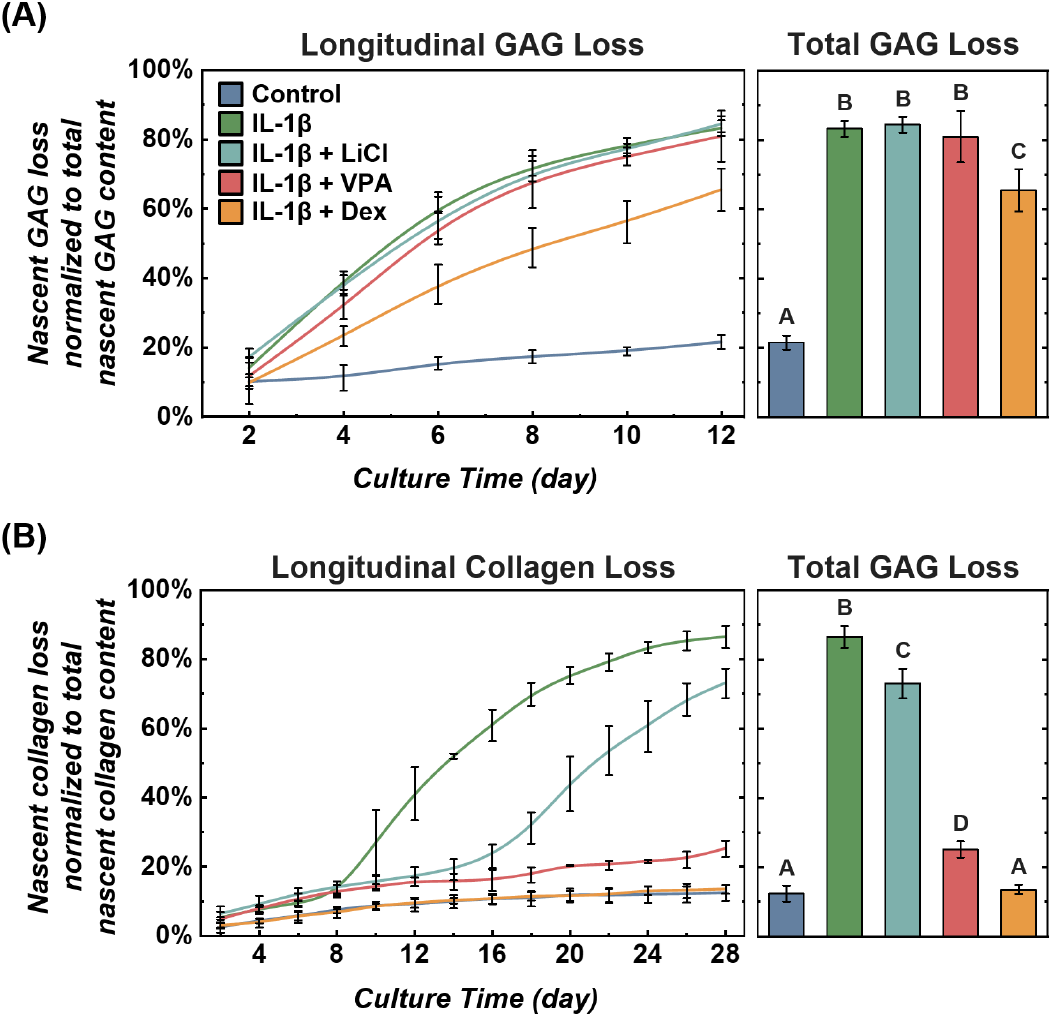
Click chemistry method for osteoarthritis drug screening. The anti-inflammatory effects of lithium chloride, valproic acid, and dexamethasone on cartilage were assessed under pro-inflammatory cytokine IL-1β stimulation (1 ng/ml) (n = 10, n = no. of samples per group). (A) Longitudinal and total loss of nascent GAG content. (B) Longitudinal and total loss of nascent collagen content. Different letters indicate significant differences between groups (p<0.05) tested via one-way ANOVA with Tukey post hoc analysis. Error bars represent ±1 SD.

The click chemistry method for tracking ECM dynamics has many potential applications in biomedical research, but the approach described here does have a few limitations. First, it incorporates modified molecular building blocks into biomacromolecules during cellular processes. This means its specificity depends on the cell or tissue type and the nature of their matrix production. For instance, it works particularly well for musculoskeletal tissues such as cartilage, given that cartilage has only two principal components, aggrecan and type II collagen. While the click chemistry method can quantify newly synthesized matrix and its degradation, it is limited to quantifying nascent ECM only. In contrast, traditional histological methods, which can reveal total ECM composition in a stable, long-term manner, remain useful. Nonetheless, the rapid, sensitive, and cost-effective quantification offered by the click chemistry method makes it a powerful tool for investigating disease mechanisms and drug effects.

## CONCLUSIONS

We developed a bioorthogonal click chemistry-based technique for rapidly and accurately quantifying ECM synthesis and degradation. This method offers significant advantages over traditional approaches. First, it is highly sensitive and accurate, enabling accurate measurement of glycan synthesis rates in as few as 10,000 MSCs every 12 hours, shortening experimental timelines from weeks to hours. This enhanced temporal resolution is especially valuable in developmental biology and regenerative medicine, where maintaining cell phenotype *in vitro* is often limited to short durations. Second, our method is cost-effective, requiring less tissue, fewer cells, and reduced labor compared to traditional techniques such as histology, chemical assays, radioisotope labeling, biochemical assays, and ELISA. It substantially decreases the overall cost and resource requirements. Third, click chemistry uniquely enables simultaneous spatial visualization and longitudinal quantification of ECM remodeling, integrating the strengths of histological staining and biochemical assays. Finally, with the continual development of new functional molecules like GalNAz and AHA, this technology can potentially label various ECM molecules synthesized by different types of cells.

Measuring nascent GAG and collagen content is essential in tissue engineering and regenerative medicine research. The click chemistry method provides a powerful platform for optimizing stem cell niches and refining the combination of cells, scaffolds, nutrients, and mechanical stimuli for artificial tissue fabrication. In addition, it holds potential for personalized medicine by enabling rapid, accurate, and cost-effective evaluations of ECM turnover in small clinical samples, such as tumor biopsies, informing therapeutic decisions and drug dosage optimization in oncology. Furthermore, the method’s speed, sensitivity, and cost-efficiency make it suited for high-throughput drug screening, particularly for conditions characterized by ECM remodeling, such as fibrosis and osteoarthritis. The advantages and potential applications of our click chemistry methods are summarized in Fig. 6.

**Figure 6.**
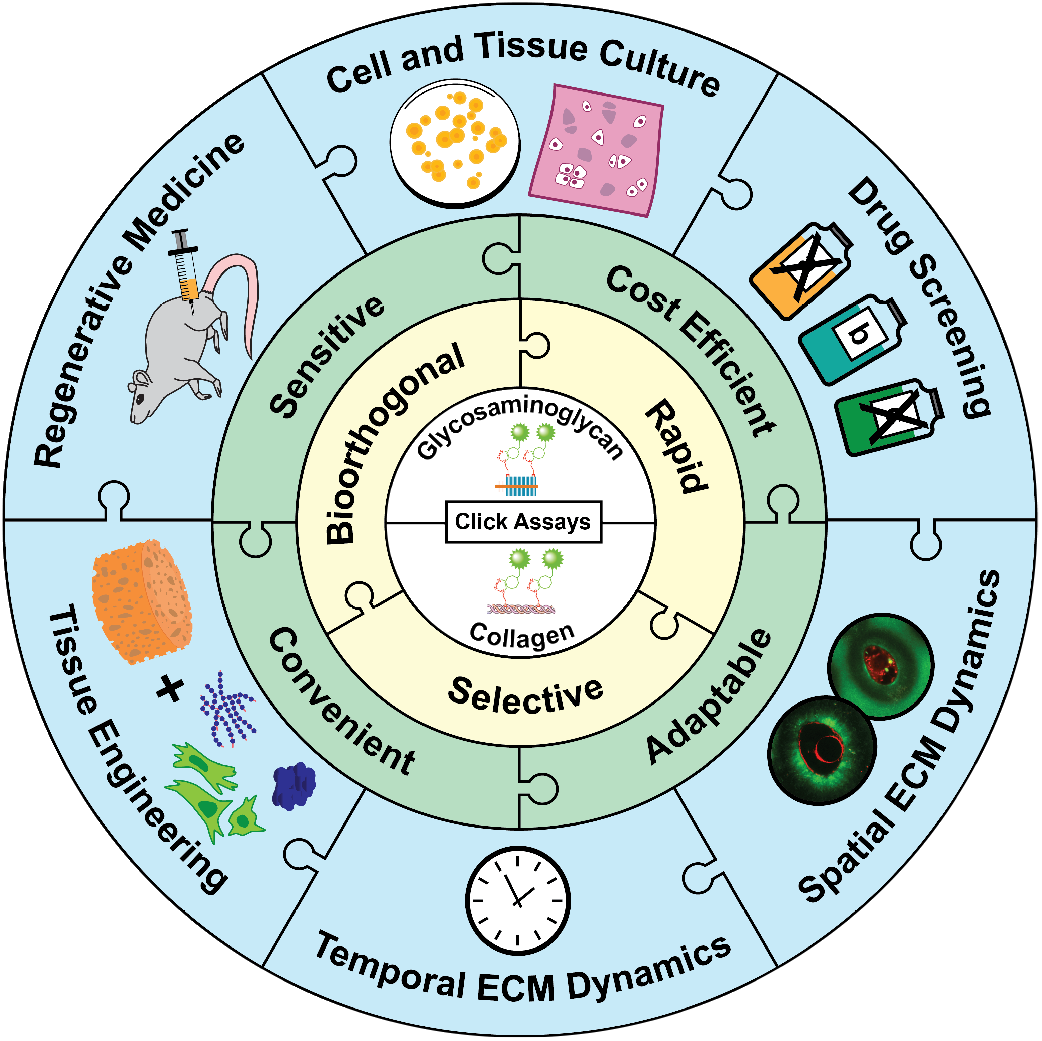
Advantages and applications of the click chemistry methods for monitoring extracellular matrix (ECM) turnover.

## MATERIALS AND METHODS

### Tissue and Cell Source

Fresh calf knee joints (1-2 months old) of mixed left or right side and sex were obtained from a local abattoir (Green Village, NJ). Cylindrical cartilage samples (diameter = 3 mm, thickness = 2 mm, tissue wet weight ≈ 15 mg) were harvested from the central, load-bearing region of the femoral condyles using a biopsy punch and custom-designed cutting tool.^21^ Cartilage samples were harvested from at least three joints for each experiment. Cartilage samples were cultured in chondrogenic medium (DMEM supplemented with 1% ITS, 0.9 mM sodium pyruvate, 50 μg/mL L-proline, and 50 μg/mL ascorbate 2-phosphate) at 37 °C, 5% CO_2_ for 48 hours before any further experiments.^21,64^

Human mesenchymal stem cells (hMSCs) and breast cancer cells (SKBR3) were used for monolayer cell culture studies. Bone marrow hMSCs were cultured in chondrogenic differentiation medium (Lonza, Walkersville, MD) supplemented with 10 ng/ml recombinant human TGF-β3 (R&D Systems, Minneapolis, MN) and expanded for use at passage 7. Cell suspensions containing 10,000 cells were seeded in 96-well plates for click chemistry labeling. SKBR3 breast cancer cells (ATCC HTB-30™, Manassas, VA) were cultured in McCoy’s 5A Medium (ATCC) and 10% FBS (Neuromics, Edina, MN) to passage 3. SKBR3 cells were seeded in 48-well plates (10,000 cells/well) containing thin polyacrylamide films of different stiffness (6, 12, and 22 kPa) to evaluate cell responses to substrate stiffness.^65^ In all experiments, cells were allowed to attach overnight before click chemistry labeling.

### Click Chemistry Labeling

To label newly synthesized GAG, 30 μM N-azidoacetylgalactosamine-tetraacylated (GalNAz, Vector Laboratories, Newark, CA) of sufficient amount was supplemented in cell/tissue culture medium for 24 hours unless otherwise specified. To label newly synthesized collagen, samples first underwent a 4-hour methionine starvation in depletion medium (DMEM-LM supplemented with 105 μg/ml Leucine, 1% ITS, 0.9 mM sodium pyruvate, 50 μg/mL L-proline, and 50 μg/mL ascorbate 2-phosphate). Samples were cultured in depletion medium supplemented with 30 μM L-Azidohomoalanine (AHA, Vector Laboratories) for 24 hours unless otherwise specified. After metabolic labeling, samples were washed thoroughly to remove excess GalNAz or AHA. A dibenzocyclooctyne (DBCO)-activated fluorescent dye, AZDye™ 488 DBCO (Vector Laboratories), was then conjugated on the azide groups of the newly synthesized ECM molecules via a copper-free click chemistry (SPAAC) reaction. Cartilage samples were submerged in 30 μM AZDye™ 488 DBCO solution for 4 hours, allowing sufficient time for the fluorophores to diffuse throughout the tissue. Monolayer cells were only dye-labeled for 30 minutes, as the click reaction happens within minutes.^66^ After fluorescent labeling, the tissue or cells were washed thoroughly in a culture medium without phenol red to remove the excess dye. No unexpected or unusually high safety hazards were encountered during the click chemistry experiments.

### Confocal Imaging and Analysis

For confocal imaging, hMSCs and SKBR3 cells were seeded in chambered coverslips and allowed to attach overnight. Cells were metabolically labeled for 48 hours with GalNAz or AHA, followed by dye labeling for 30 minutes with AZ488 DBCO. Cells were counterstained with CellTracker™ Red CMTPX Dye (1X concentration; ThermoFisher, Waltham, MA) for 45 minutes and nuclei were counterstained with Hoechst 33342 (1 μg/ml; ThermoFisher) for 10 minutes. Cartilage samples were bisected into two hemicylinders after click chemistry labeling. Chondrocytes in the samples were counterstained with CellTracker™ Red CMTPX Dye (2X concentration) for 5 hours followed by thorough washing. Fluorescent images were taken with a 20X, 40X, or 60X objective on a Zeiss LSM880 multiphoton confocal microscope. Cartilage samples were imaged at a focal plane ∼50 μm below the cross-section surface to avoid cells damaged during the preceding sample preparation. All fluorescent images were processed in ImageJ.^67^

### Histology

Cartilage samples were fixed for 24 hours in 10% formalin and then paraffin embedded. Samples were cut into 5 μm thick sections and stained with Safranin O/Fast Green to assess GAG distribution or Picrosirius Red to assess collagen distribution.

### Quantification of ECM Synthesis and Loss

Cartilage samples were digested with papain (125 µg/ml) for 48 hours to quantify ECM synthesis following click chemistry labeling (n = 10 samples per group).^20^ The digested solution was diluted if necessary, and then the fluorescent intensity was read on a plate reader (Gemini EM; Molecular Devices, San Jose, Ca). The fluorescent reading was normalized to the sample wet weight obtained immediately after tissue harvest and further normalized to the corresponding control samples (n = 10, n = no. samples per group). The amount of newly synthesized ECM in each sample was proportional to the normalized fluorescent intensity. Synthesis by monolayer cells was quantified with a similar protocol. Fluorescent intensity of each hMSCs group was normalized to the first 12-hour timepoint, and to the 6 kPa group for SKBR3 cells (n = 10, n = no. of replicates) To track the longitudinal loss of ECM in cartilage, click chemistry labeled samples were continuously cultured with proinflammatory cytokine IL-1β (1ng/ml, RBOIL1BI, ThermoFisher) to promote ECM degradation. Samples were cultured for either 10 days (to track GalNAz-labeled GAG loss) or 28 days (to track AHA-labeled collagen loss), per the differing GAG and collagen loss kinetics reported in the literature (n = 10, n = no. samples per group).^58^ The click-labeled ECM degraded and diffused out of the cartilage sample. The culture medium was replaced every other day, and the fluorescence of the old medium was quantified on a plate reader. At the end of the cartilage culture, samples were digested with papain as described above, and the fluorescence of the digestion solution was obtained. The total amount of labeled GAG or collagen was represented by the sum of the medium longitudinal readings and the digestion solution, which was subsequently used to calculate the longitudinal ECM loss.

### Comparative Chemical Assays for GAG and Collagen Content

Results from the click-chemistry method were compared with traditional chemical assays. GAG content in culture medium and cartilage was measured with a dimethylmethylene blue (DMMB) dye-binding assay, with chondroitin-4-sulfate as a standard.^57^ All chemicals were obtained from the Blyscan™ Glycosaminoglycan Assay kit (Biocolor Ltd, UK). The collagen content was determined by colorimetric hydroxyproline quantification method via dimethylaminobenzaldehyde (DMAB) and chloramine T assay. The hydroxyproline-collagen ratio was assumed as 1:7.5 for calf cartilage.^4^

### Statistical Analysis

Statistical analyses were conducted in R (R Version 4.1.1, R Core Team 2021).^68^ All data were presented as means ± standard deviation. Before analysis, normality and homoscedasticity assumptions were assessed using Shapiro-Wilks and Q-Q plots, respectively. A *p* value of less than 0.05 was considered statistically significant. To assess the correlation between substrate stiffness and ECM synthesis in SKBR3 cells, a simple linear regression was conducted. All other synthesis experiments were assessed using one-way ANOVA with Tukey post hoc tests to compare all means and adjust for family-wise type I errors. The cumulative loss of GAG and collagen was compared using one-way ANOVA with Tukey post hoc tests either at each of the tested time points (IL-1β dosage study) or only at the final time point to compare total GAG loss (drug screening study). To evaluate the correlation between the click chemistry and traditional chemical assays, simple linear regression on the individual non-cumulative readings at all time points was performed.

## Supplementary Material

Methionine content of abundant cartilage matrix proteins [Table S1], GAG and collagen temporal loss profiles of individual samples [Figure S1] (DOC).

## Acknowledgments

This work was financially supported by the Department of Defense (DOD) Grant W81XWH-13-1-0148 (to XLL), National Institutes of Health (NIH) Grant R01AR074472 (to XLL), P20GM139760, and R01AR074490 (to LH), as well as National Science Foundation (NSF) Grant CMMI-1751898 (to LH). AP was supported by NSF Graduate Research Fellowship Program (GRFP) and the Helwig Fellowship in Mechanical Engineering at the University of Delaware.

