## Supplemental Table 1 and Figure 1 for "Click chemistry-based quantification of extracellular matrix turnover for drug screening and regenerative medicine"

This supporting information contains:

Table S1. Methionine Content of ECM Proteins S2
Figure S1. Illustration of ECM loss profiles for individual cartilage samples tracked by click chemistry and traditional chemical assays. S3

**Table S1:** Methionine content of select common ECM proteins in bovine articular cartilage. Retrieved from UniProtKB/Swiss-Prot.[1]

| UniProtID | Protein Name | Gene | # of Amino Acids | # of Methionines | **Methionine Content (%)** |
|---|---|---|---|---|---|
| P02459 | Collagen alpha-1(II) chain | COL2A1 | 1,487 | 20 | **1.34** |
| C0HLN2 | Collagen alpha-2(IX) chain | COL9A2 | 688 | 10 | **1.45** |
| Q28083 | Collagen alpha-1(XI) chain | COL11A1 | 911 | 10 | **1.10** |
| Q32S24 | Collagen alpha-2(XI) chain | COL11A2 | 1,736 | 19 | **1.09** |
| P13608 | Aggrecan core protein | ACAN | 2,364 | 10 | **0.42** |
| P35445 | Cartilage oligomeric matrix protein | COMP | 756 | 11 | **1.45** |

[1] The UniProt Consortium. UniProt: The Universal Protein Knowledgebase in 2023. *Nucleic Acids Res.* **2023**, *51* (D1), D523–D531. https://doi.org/10.1093/nar/gkac1052.

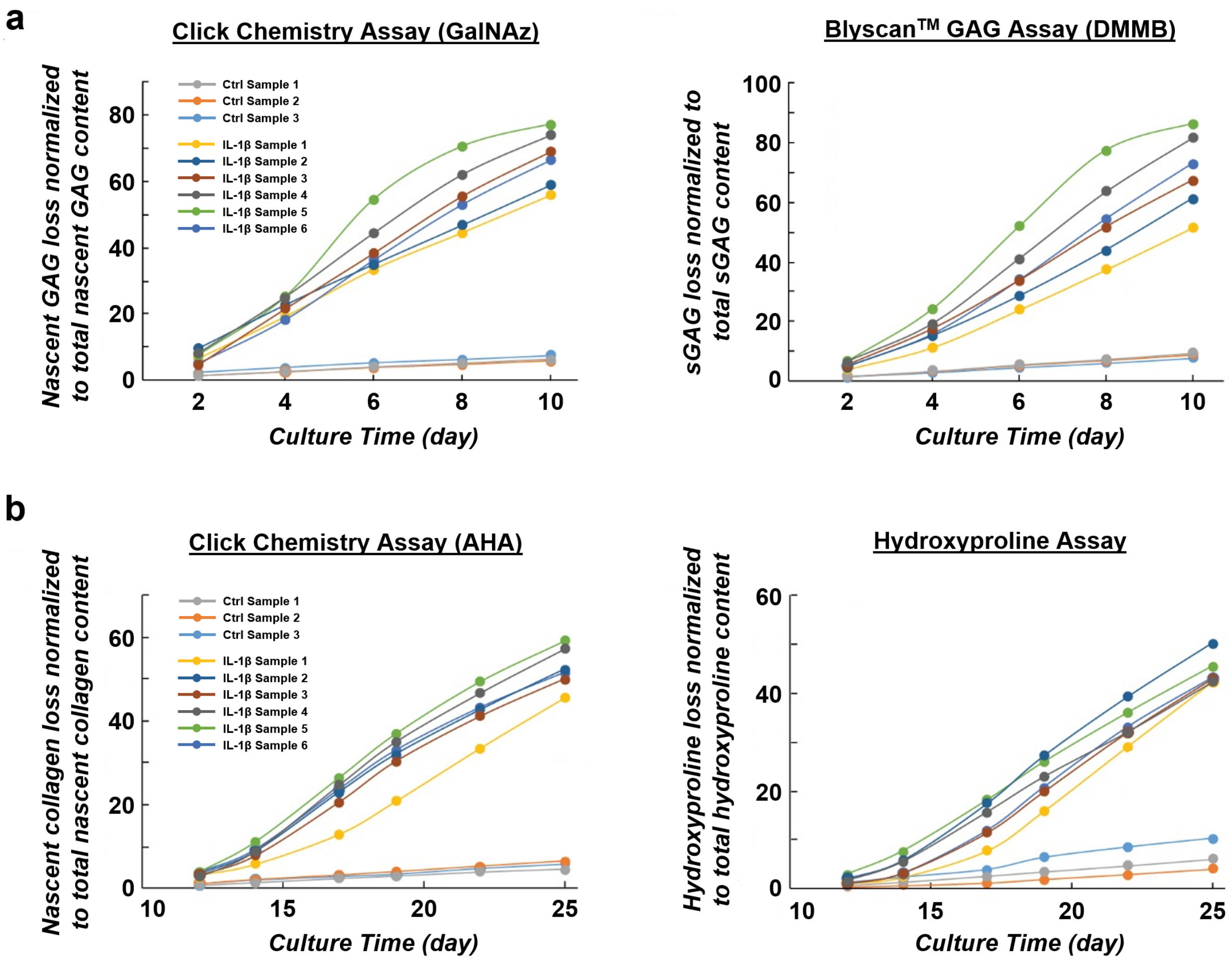


**Figure S1. Illustration of ECM loss profiles for individual cartilage samples tracked by click chemistry and traditional chemical assays.** Illustrated here are the longitudinal ECM loss profiles for individual cartilage samples (n=3 ctrl, n=6 IL-1β). (a) The loss of glycosaminoglycans (GAG) tracked by click chemistry using a GalNAz tag and the traditional dimethylmethylene blue (DMMB) assay for sulfated GAGs. The loss of GAG into the culture media from cartilage samples exposed to the pro-inflammatory cytokine interleukin 1-beta (IL-1β; 1ng/ml) was tracked for 10 days. (b) The loss of collagen tracked by click chemistry using an AHA tag and the traditional hydroxyproline assay. The loss of collagen into the culture media from cartilage samples exposed to the pro-inflammatory cytokine IL-1β (1 ng/ml) was tracked for 25 days.
